# Deciphering lipidomics landscape of urine extracellular vesicles in alcohol use disorder patients

**DOI:** 10.1101/2025.05.27.656263

**Authors:** Blanca Martín-Urdiales, Carla Perpiñá-Clérigues, Susana Mellado, Maura Rojas-Pirela, María-Lourdes Aguilar Sánchez, David Puertas-Miranda, Francisco García-García, Miguel Marcos, María Pascual

**Affiliations:** Department of Physiology, School of Medicine and Dentistry, University of Valencia, 46010 Valencia, Spain; Computational Biomedicine Laboratory, Príncipe Felipe Research Center, 46012 Valencia, Spain; Department of Internal Medicine, University Hospital of Salamanca, 37007, Salamanca, Spain; Institute of Biomedical Research of Salamanca (IBSAL), 37007 Salamanca, Spain; Primary Care Addiction Research Network (RIAPAd), Instituto de Salud Carlos III, 28029, Madrid, Spain; Department of Medicine, University of Salamanca, 37001, Salamanca, Spain; Psychiatry Service, University Hospital of Salamanca, 37007, Salamanca, Spain

**Keywords:** Lipidomics, lipid network, extracellular vesicles, urine, alcohol use disorder

## Abstract

Urine extracellular vesicles (EVs) have garnered increasing interest in recent years as, in combination with their lipid content, they represent a potential source of non-invasive biomarkers. Alcohol use disorder (AUD) is one of the most common psychiatric disorders and a significant contributor to morbimortality. Thus, there is growing interest in assessing previous alcohol consumption and evaluating the severity of liver or brain injury, particularly via non-invasive methods. Here, we employed a novel approach based on urine EVs and a highly sensitive lipidomics strategy to characterize lipid species in male AUD patients, as well as to evaluate the differential functional roles and enzymatic activity networks of urine EV lipids. Our results show, for the first time, that the lipidomic profiling of urine EVs in AUD males is characterized by an enrichment of fatty acyls and glycerophospholipids, with FA 22:0 emerging as a potential biomarker. Notably, increased acyl chain saturation and elevated long-chain fatty acids (22-24 carbons) suggest links to AUD-associated inflammation, cancer, and metabolic dysfunction. These findings support an improved understanding of the urine EV lipidome, which could contribute to the identification of novel lipid targets and the discovery of non-invasive biomarkers in AUD.

## Introduction

Extracellular vesicles (EVs) are released by all types of living cells and they play important roles as intercellular mediators in numerous physiological and pathological processes (Pascual et al., 2020; Shao et al., 2018). Due to their composition of proteins, nucleic acids, and lipids, EVs have been recognized as promising biomarker targets in a variety of diseases (Shao et al., 2018). Urine EVs, in particular, have garnered interest as a potential source of non-invasive biomarkers, as they can reflect molecular events related to pathological alterations not only in the urinary system but also with other diseases, such as neurodegenerative disorders and cancer (Erdbrügger et al., 2021; Fraser et al., 2016; Oeyen et al., 2019; Yasui et al., 2024). Since the cargo of EVs is thought to reflect the cell-type of origin, analyzing the urinary EV cargo can may yield biological markers for the diagnosis or progression of various pathological condition (Erdbrügger et al., 2021; Oeyen et al., 2019).

Whereas proteins and microRNA in plasma EVs have been widely analyzed (Zhu et al., 2022), the lipid content of urine EVs remains underexplored and represents a promising source of non-invasive biomarkers (Lyu et al., 2024; Oeyen et al., 2019; Zhu et al., 2022). Mass spectrometry-based lipidomics, combined with dedicated computational tools (Rose et al., 2023), represents a powerful approach for identifying and quantifying lipids in body fluids (Han & Gross, 2022), including urine (Lyu et al., 2024; Oeyen et al., 2019; Zhu et al., 2022). Understanding how pathological conditions alter lipids and their impact on cellular processes is a critical challenge that could provide new insights into disease mechanisms (Han & Gross, 2022). In this context, novel bioinformatic approaches such as the Lipid Network Explorer (LINEX^2^) (Rose et al., 2023), which integrates lipid classes and lipid-metabolizing enzyme activity, offer a comprehensively method for identifying novel lipid targets and discovering clinical biomarkers (Lv et al., 2018; Perpiñá-Clérigues et al., 2024, 2025).

Alcohol use disorders (AUD) represents a heterogeneous spectrum of clinical manifestations, with the liver and brain being the organs most affected by excessive alcohol consumption (Wolstenholme et al., 2024). AUD is estimated to cause approximately three million deaths globally each year and is a significant contributor to morbimortality (Ayares et al., 2022). Alcohol-induced health consequences to include alcohol-associated liver disease, hepatocellular carcinoma, cardiac disease, and psychiatric disorders (Ayares et al., 2022). There is growing interest in developing tools to assess prior alcohol consumption and evaluate the severity of liver injury, particularly through non-invasive methods.

Our previous studies identified the lipidomic fingerprint in plasma EVs from ethanol-intoxicated adolescents (Perpiñá-Clérigues et al., 2023) and AUD patients (Perpiñá-Clérigues et al., 2024). Advancing previous research, the present study employs an innovative approach using urine EVs and a highly sensitive lipidomics strategy to characterize lipid species in male AUD patients and evaluate the functional roles and enzymatic activity networks of urine EV lipids. We demonstrate, for the first time, that the lipidomic profile of urine EVs in AUD males is characterized by an enrichment of fatty acyls and glycerophospholipids, with FA 22:0 as a potential biomarker. Notably, increased acyl chain saturation and elevated long-chain fatty acids (22-24 carbons) suggest links to AUD-associated inflammation, cancer, and metabolic dysfunction.

## Material and methods

### Human subjects

Seven male AUD patients, diagnosed according to DSM-5 criteria, were recruited for this study through Alcoholism Unit at the University Hospital of Salamanca (Spain). The median age of the AUD group was 43 years. All patients reported active alcohol consumption ≥ 100 g of ethanol per day prior to study inclusion. Laboratory results showed normal prothrombin time, hemoglobin concentration, and serum albumin levels. All patients tested negative for hepatitis B surface antigen and hepatitis C antibodies. None of the participants had acute or chronic conditions that could interfere with the study results, nor were they polydrug users. Advanced liver disease was excluded based on clinical examination, laboratory analysis, and ultrasonography. Patients with physical signs of chronic liver disease (e.g., cutaneous stigmata, hepatosplenomegaly, gynecomastia, testicular atrophy, and/or muscle wasting), ultrasonographic abnormalities beyond hepatic steatosis, or liver transaminase levels exceeding 2–3 times the reference range were excluded (Table S1). Seven healthy male volunteers, who reported consuming < 15 g of ethanol per day, were included as controls. Their median age was 47 years. All participants provided written informed consent, and the study was approved by the Ethics Committee of the University Hospital of Salamanca, Spain (Project ID: PI 2023 07 1389). Urine samples were collected in sterile, additive-free 50 mL polypropylene tubes and stored at-80 °C until further processing.

### EV isolation from human urine

To remove cell debris, urine samples were centrifuged at 300 x g for 10 min. The supernatant was then centrifuged at 2,000 x g for 10 min, and the resulting supernatant was transferred to clean tubes and ultracentrifuged at 100,000 x g for 2 h at 4 ºC. The resulting EV pellet was washed with phosphate-buffered saline (PBS) and centrifuged again at 100,000 x g for 1 h. The final pellet was used either for lipid extraction or for EV characterization.

### Urine EV characterization by transmission electron microscopy, Western blot, and nanoparticle tracking analysis

Freshly isolated EVs were resuspended in PBS and characterized using transmission electron microscopy, Western blot, and nanoparticle tracking analysis., following previously described protocols (Ibáñez et al., 2019). For transmission electron microscopy, EVs were examined with an FEI Tecnai G2 Spirit microscope (FEI Europe, Eindhoven, The Netherlands) equipped with a Morada digital camera (Olympus Soft Image Solutions GmbH, Münster, Germany). Western blot analysis employed antibodies against CD9, CD63, CD81, and calnexin (Santa Cruz Biotechnology, USA). Representative blots for each marker are shown in Figure S1. EV size and distribution and concentration were assessed using a NanoSight NS300 system (Malvern Panalytical., Malvern, UK).

### Lipid extraction

Lipids from EVs were extracted from equal volumes (21 mL/sample) of urine using a modified Folch method. The lipid-containing lower phase was transferred to clean tubes, dried under nitrogen, and stored at-80 °C. Dried lipid samples were resuspended in isopropanol for subsequent analysis by liquid chromatography-tandem mass spectrometry (LC-MS/MS) in both positive and negative ionization modes.

### Liquid chromatography-MS/MS analysis

Lipid extracts were analyzed using a quadrupole time-of-flight (QTOF) mass spectrometer operated in auto MS/MS mode. A pooled sample representing all fourteen biological replicates (two groups × seven samples) was analyzed iteratively. Detailed experimental methods for liquid chromatography (LC) and auto MS/MS were performed as previously described (Sartain et al., 2019, 2020), with minor modifications. Lipid separation was achieved using an Agilent 1290 Infinity LC system coupled to the 6550 Accurate-Mass QTOF instrument (Agilent Technologies, Santa Clara, CA, USA) equipped with an electrospray interface (Jet Stream Technology, Agilent Technologies) operating in either positive-ion mode (3500 V) or negative-ion mode (3000 V) and high sensitivity mode. Optimal conditions for the electrospray interface included a gas temperature of 200 °C, drying gas flow of 12 L/min, nebulizer pressure of 50 psi, sheath gas temperature of 300 °C, and sheath gas flow of 12 L/min. Lipids were separated on an Infinity Lab Poroshell 120 EC-C18 column (3.0 × 100 mm, 2.7 μm; Agilent Technologies). The mobile phase consisted of solvent A (10 mM ammonium acetate, 0.2 mM ammonium fluoride in 9:1 water/methanol) and solvent B (10 mM ammonium acetate, 0.2 mM ammonium fluoride in 2:3:5 acetonitrile/methanol/isopropanol) using the following gradient: 0 min 70% B, 1 min 70% B, 3.5 min 86% B, 10 min 86% B, 11 min 100% B, 17 min 100% B. The separation was performed at 50 °C with a constant flow rate of 0.6 mL/min. The injection volume was 5 μL in both positive and negative modes.

Data acquisition was performed using Agilent MassHunter Workstation Software B.10.1 in auto MS/MS mode, selecting the top three ions (charge states, 1-2) within 300–1700 m/z range exceeding 5,000 counts and 0.001% were selected for analysis. The quadrupole was set to a “narrow” resolution (1.3 m/z), and MS/MS spectra (50–1700 m/z) were acquired up to 25,000 total counts or 333 ms. Mass accuracy was maintained using continuous internal calibration using m/z 121.050873 and m/z 922.009798 signals for positive mode and m/z 119.03632 and m/z 980.016375 signals for negative mode. Additionally, all ions MS/MS (Agilent Technologies, 2013) data were acquired individually with a scan rate of 3 spectra/s and collision energies of 0, 10, 20, and 40 eV.

### Lipid annotator database

Pooled MS/MS data (five sets of five iterations) were analyzed using Lipid Annotator software 1 (Koelmel et al., 2020) which applies Bayesian scoring and non-negative least-squares to match spectra against a modified LipidBlast library developed by Kind et al., 2013 (Tsugawa et al., 2015).

The software used was Agilent MassHunter Lipid Annotator v1.0, with precursor ion selection limited to [M+H]+ and [M+NH4]+ in positive ion mode and [M-H]– and [M+HAc-H]-in negative mode. The Agilent MassHunter Personal Compound Database and Library (PCDL) Manager vB.08 SP1 was used to manage annotations.

### Lipid identification

The custom lipid PCDL databases were used for batch-targeted feature extraction in Agilent MassHunter Profinder (v10.0), processing fourteen all-ions MS/MS data files. We employed the provided “Profinder - Lipids.m” method with modifications as described by Sartain et al., 2020. Feature identification was performed using the Find by Formula (FbF) algorithm, adapted for all-ions MS/MS analysis. The algorithm first compares mass peaks in the low-energy channel against the PCDL database entries with matching m/z values, generating a list of putative identifications. These candidates were then verified by matching fragment ions from the PCDL against detected ions in the high-energy channel.

For each candidate, precursor and product ions were extracted as chromatograms and evaluated using a coelution score that incorporates abundance, peak symmetry, width, and retention time. All results were manually reviewed in MassHunter Profinder, with ambiguous features (poor peak shape or integration quality) being either manually corrected or excluded. Finally, qualified features were exported as a.cef file for downstream analysis.

### Bioinformatic analyses

Lipidomic bioinformatic analysis was performed in R software v.4.1.2 (R Development Core Team, s. f.). The experimental design and analytical workflow are illustrated in Figure 1. Comparison focused in bioinformatic analysis was performed between AUD male patients versus healthy individuals (AUD - C).

**Figure 1.**
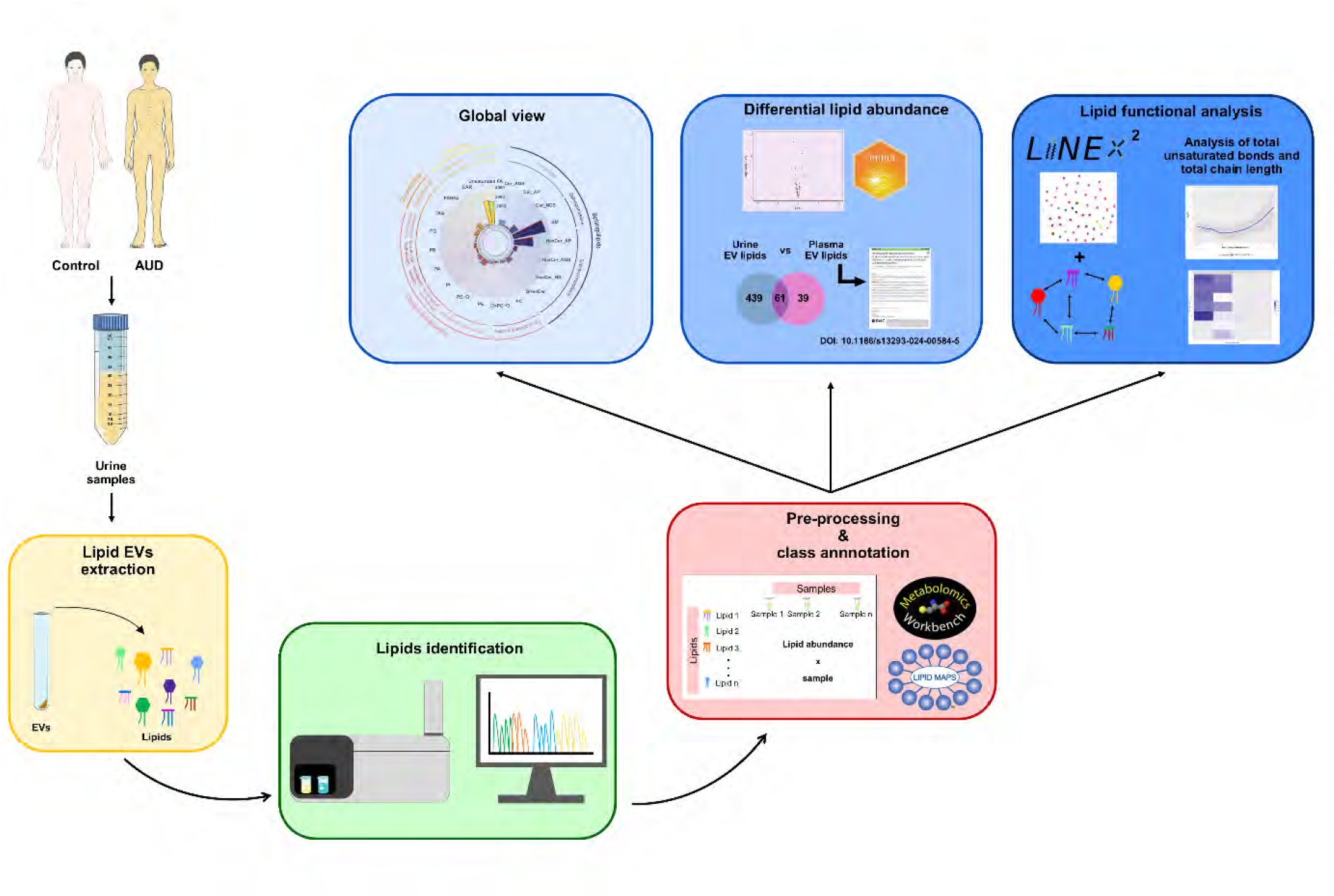
Experimental design and lipidomic workflow. Urine extracellular vesicles (EVs) were isolated from male patients with alcohol use disorder (AUD) and healthy individuals (AUD vs. C). Lipids were extracted, quantified, and identified using LC-MS/MS. After data normalization and lipid class annotation, exploratory and differential analyses were performed to assess lipid abundance. Venn diagrams illustrate overlapping lipid species between urine EVs and our previously published data from plasma EVs of males with AUD (Perpiñá-Clérigues et al., 2024). For functional analysis, the LINEX^2^ platform was employed to generate a global reaction network and a subnetwork highlighting the most significant average substrate-product change. Additionally, the total number of unsaturated bonds and total chain length (carbon number) was analyzed in urine EVs lipid species from AUD patients and controls.

### Data preprocessing

Data preprocessing involved entity filtering, lipid abundance matrix normalization, and exploratory analyses. Raw data from MassHunter Qualitative (.cef files) were imported into Mass Profiler Professional (Agilent Technologies) for statistical processing. Entities were filtered by frequency, retaining only those consistently detected in all replicates of at least one experimental group.

The data were normalized using a 75th percentile shift algorithm and baselined to the median of all samples. Normalized datasets were labeled by ionization mode (negative/positive) and consolidated into a single data frame. Exploratory analyses included hierarchical clustering, principal component analysis (PCA), and sample/lipid-specific box-and-whisker plots to identify abundance patterns and detect potential batch effects.

Samples exhibiting anomalous behavior (defined as values exceeding 1.5 × interquartile range (IQR) below Q1 or above Q3) were excluded to minimize batch effects that could significantly impact differential abundance analysis.

### Differential lipid abundance

Differential abundance was assessed using the limma R package (Ritchie et al., 2015), with p-values adjusted by Benjamini & Hochberg (BH) procedure (Benjamini & Hochberg, 1995). The log2 fold change (LFC) statistic indicated whether lipid abundance was greater in AUD patients (positive LFC) or controls (negative LFC). Shared biomarkers between urinary and plasma EVs in AUD were identified using previously published data (Perpiñá-Clérigues et al., 2024).

### Class annotation

Lipid class annotation was performed using the RefMet (Fahy & Subramaniam, 2020) and LIPID MAPS database (Sud et al., 2007). Abbreviations are defined in Table S2. Lipids were ranked by p-value and direction of change (Table S4).

### Lipid network

The Lipid Network Explorer platform (LINEX^2^, https://exbio.wzw.tum.de/linex/) was used to analyze lipid metabolic networks and investigate sex-specific lipid dysregulation in AUD patients (Rose et al., 2023). Lipids were analyzed as molecular species regardless of retention time or ion mode. Lipid nomenclature was standardized using MetaboAnalyst 5.0 platform (Pang et al., 2021) and the LipidLynxX Converter tool (http://www.lipidmaps.org/lipidlynxx/converter/) (Ni & Fedorova, 2020), and manually verified. LINEX^2^ analysis provided global lipid species networks based on reaction types and Spearman correlations. Network enrichment analysis identified subgraphs with the most significant substrate-product changes, proposing enzymatic dysregulation via a greedy local search algorithm.

### Lipid structure analysis

The total number of acyl carbon atoms (chain length) and unsaturated bonds was determined using the lipidr R package (Mohamed et al., 2020). This package extracts structural features by parsing lipid names, including the combined fatty acyl chain length and total number of double bonds per lipid species. Distributional changes in these features between AUD and control conditions were then analyzed using built-in lipidr functions. Specifically, locally weighted regression (LOESS) was applied to model the relationship between log2 fold-change (log2FC) and either chain length or degree of unsaturation. For univariate analysis, significantly regulated lipid species were identified using a moderated t-statistic implemented in lipidr.

### Web platform

The interactive web platform (https://bioinfouvcipf.shinyapps.io/LipUrEVs-AUD/) provides comprehensive documentation of the computational methodologies employed in this study. This resource enables users to explore statistical results from all analyses and identify lipid profiles of interest. As an open-access tool, it aims to facilitate data sharing, support innovative research, and promote the discovery of novel findings in the field.

## Results

### Lipidomics data exploration of urine EVs isolated from male patients with AUD

Before analyzing the lipid profile of urine EVs in healthy individuals and AUD patients, we characterized the isolated EVs using electron microscopy, Western blot, and nanoparticle tracking analysis (Fig. 2). TEM analysis confirmed that the nano-sized particles exhibited typical exosomal morphology and size (∼100 nm in diameter; Fig. 2A). These EVs expressed the exosomal markers CD63, CD9, and CD81 (tetraspanin proteins) and showed no cytosolic contamination, as evidenced by the absence of calnexin (Fig. 2B). Nanoparticle tracking analysis further supported these findings, with a peak within the 100-200 nm range, consistent with EVs (Fig. 2C).

**Figure 2.**
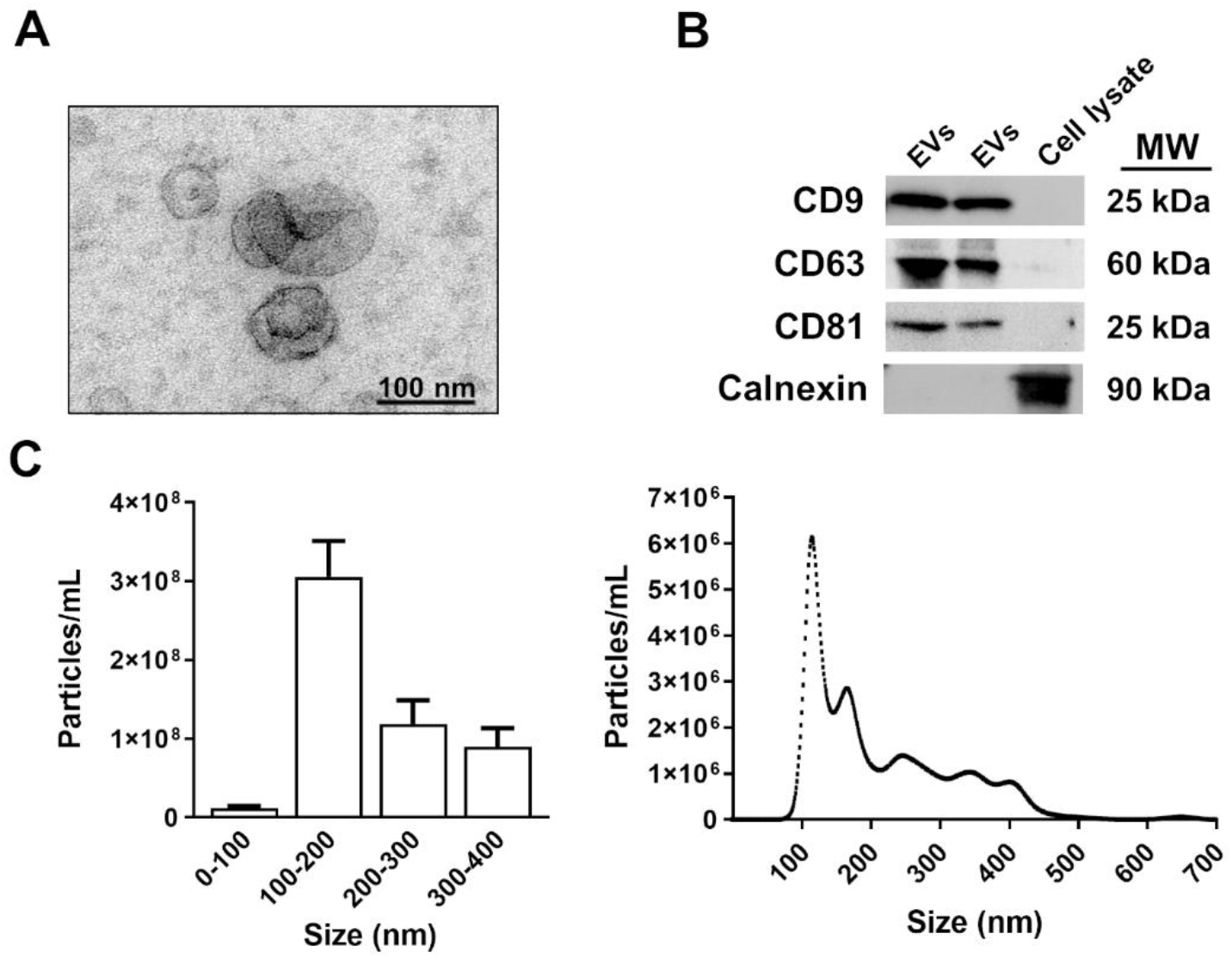
Characterization of urine extracellular vesicles (EVs). **(A)** Transmission electron microscopy image of isolated human urine EVs. **(B)** Immunoblot analysis of EV markers (CD9, CD63, and CD81) in urine EVs and astroglial cell lysates (positive control for calnexin). Calnexin expression was assessed to rule out cytosolic protein contamination in EV isolates. A representative blot is shown for each target. **(C)** Nanoparticle track analysis of urine EVs: size distribution (left) and particle concentration (right).

To analyze the lipidomic profiles of urine EVs, we identified 105 lipid species using the RefMet and LIPID MAPS databases, spanning the superclasses sphingolipids, glycerophospholipids, fatty acyls, and glycerolipids. While most subclasses showed low abundance, we observed an enrichment of unsaturated FA, HexCer_AP, and SM in urine EVs (Fig. 3A and Table S3). The lipid abundance distribution was similar between control and AUD males (Fig. 3B). However, hierarchical clustering of EV lipid species revealed distinct profiles between the two groups, irrespective of subclass (Fig. 3C and D), suggesting that alcohol consumption contributes to sample variance (see PC1 in Fig. 3D).

**Figure 3.**
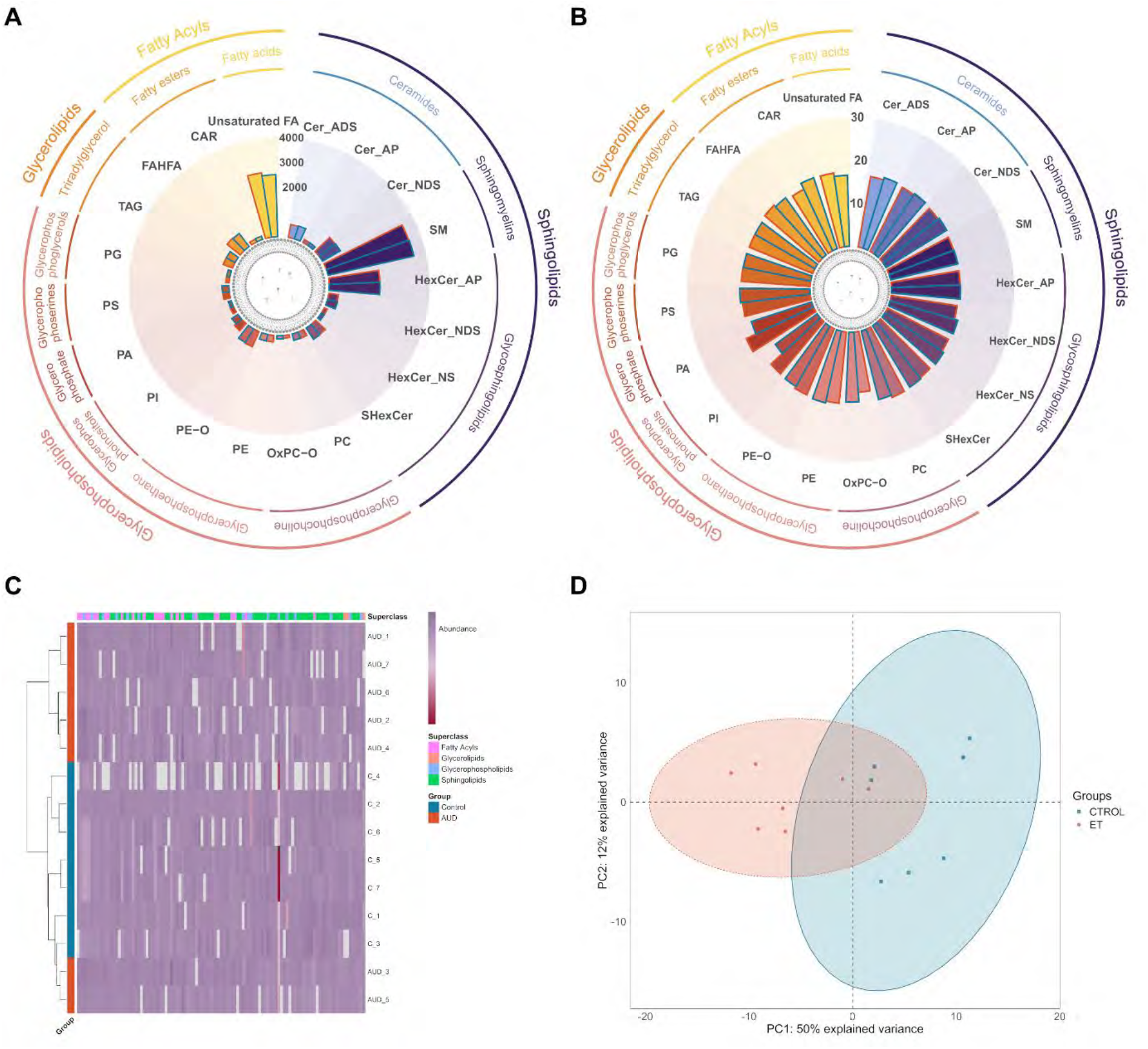
Lipid composition and distribution of lipids from urine extracellular vesicles (EVs) from male alcohol use disorder (AUD) patients and controls. **(A)** Total and **(B)** average lipid abundance (log2-transformed LC-MS/MS peak area) by lipid subclass, grouped by RefMet classification (inner line: main class; outer line: superclass). Bar border colors denote patient groups (AUD: red and control: blue). Lipid abbreviations are detailed in Table S2. **(C)** Heatmap of lipid abundance patterns (columns: lipids; rows: samples). Abundance levels are represented on a violet-maroon scale, with maroon indicating lower and violet higher abundance. **(D)** Principal component analysis (PCA) score plot revealing distinct clustering by patient group (color-coded as above).

### Analysis of lipid abundance of urine EVs isolated from male individuals with AUD

We then assessed significant variations in lipid abundance in urine EVs between AUD males and controls. Figure 4A shows that the superclasses with higher lipid abundance in AUD males compared to controls were fatty acyls and glycerophospholipids, whereas sphingolipids displayed lower lipid abundance in AUD males. In addition, fifteen lipid species displayed significant alterations (p-value ≤ 0.05) when comparing AUD males to controls (Figure 4B and Table S4). Specifically, the PI 34:1, PA 2:0_16:3, FA (22:1, 19:0, 21:0, 22:0) subclasses revealed significantly altered lipids with greater abundance in AUD males, while ACar 10:1, Cer_ADS d43:0, Cer_NDS d44:1 and SM d35:1 subclasses displayed significantly altered lipids with lower abundance in AUD males.

**Figure 4.**
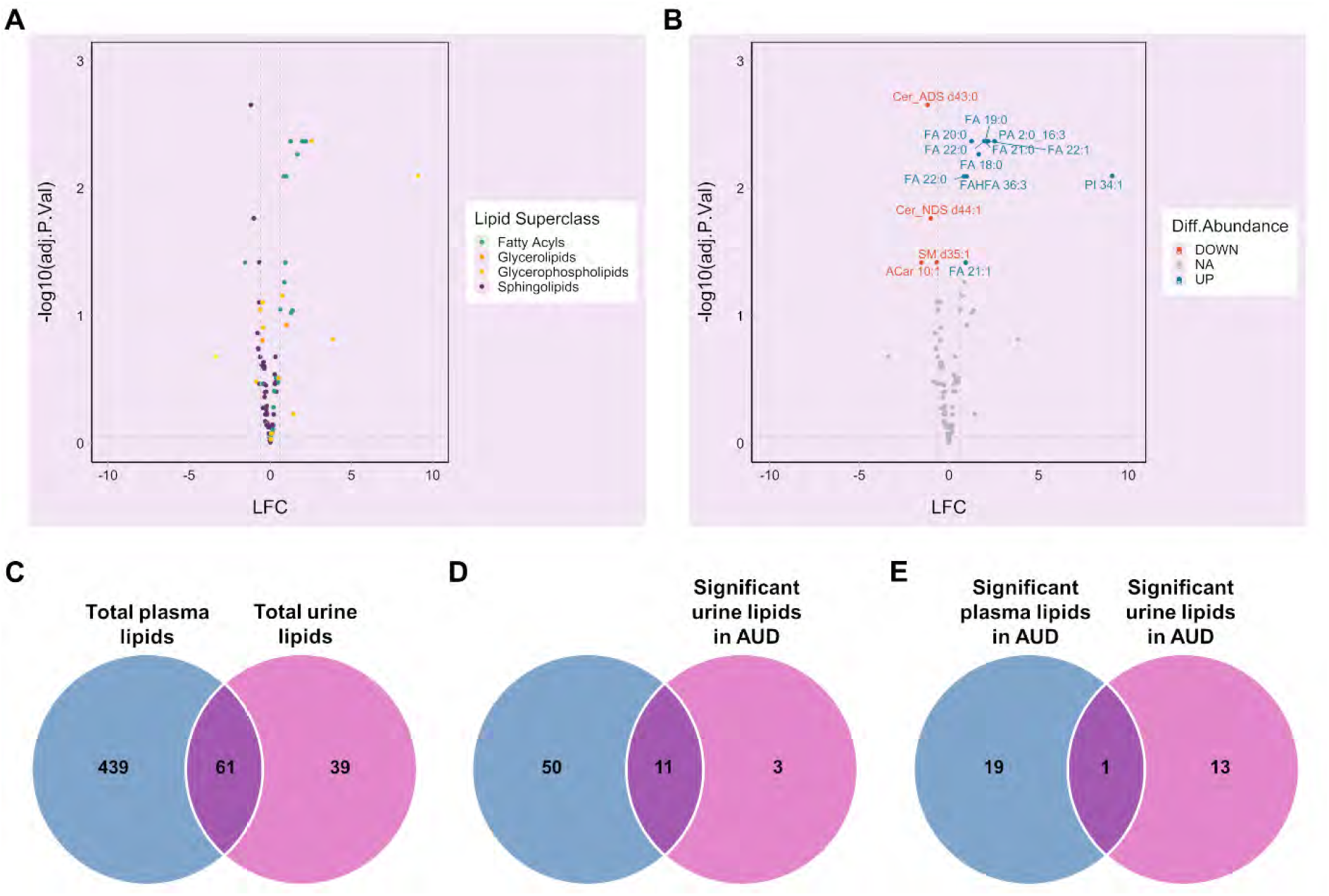
Differential lipid abundance in urine extracellular vesicles (EVs) from male alcohol use disorders (AUD) patients versus controls. **(A)** Volcano plot of all lipid species (colored by superclass) and their log2 fold changes (LFCs) in AUD. **(B)** Volcano plot highlighting significantly altered lipid subclasses (adjusted p ≤ 0.05; red: higher abundance in AUD; blue: lower; gray: nonsignificant). **(C)** Venn diagram of lipid species overlapping between urine and plasma EVs in AUD. **(D)** Venn diagram of lipids with significant abundance changes (LFC > 1, adjusted p ≤ 0.05) in urine EVs that are also detected in plasma EVs. **(E)** Venn diagram of lipids with significant abundance changes (LFC > 1, adjusted p ≤ 0.05) in urine EVs that are also detected in plasma EVs.

We then analyzed the lipidome data from urine and plasma EVs isolated from male patients with AUD and healthy individuals, using the plasma lipidome data previously published in Perpiñá-Clérigues et al., 2024. Venn diagram in Figure 4C, which compares the lipidome of urine and plasma EVs, revealed sixty-one common lipid species in both fluids. In addition, eleven lipid species that showed a significant abundance changes in urine EVs were also present in plasma EVs (Fig. 4D). However, when comparing the significantly altered lipid species in both fluids, we found that FA 20:0 was upregulated in urine and plasma EVs from AUD males compared to controls (Fig. 4E).

### Analysis of the lipid network / Lipid functional analysis of urine EVs isolated from male AUD patients

To derive biological insights from lipidomics data, we utilized LINEX^2^. Figure 5A displays the global network of lipid species, illustrating qualitative relationships between species based on predefined reaction types. Most reactions relate to SM and Cer modification/removal (orange and blue edges). Notably, fatty acyls are not detected in this functional analysis since LINEX^2^ is not configured to analyze free fatty acyls. Figure 5B depicts qualitative associations regarding alterations in lipid levels between AUD and control males. The size of the spherical nodes represents higher and lower lipid abundance based on the –log10(FDR). Figure 5C highlights the most dysregulated subnetwork between AUD males and control individuals. While the spherical nodes represent lipid species, the triangular nodes represent reaction type. The resulting enzymatic dysregulation reactions include only SM and Cer in these patients, consistent with data obtained from plasma EVs of AUD males (Perpiñá-Clérigues et al., 2024). This suggests that these lipid species, which are dysregulated by different biochemical reactions, are detected in EVs from both biofluids.

**Figure 5.**
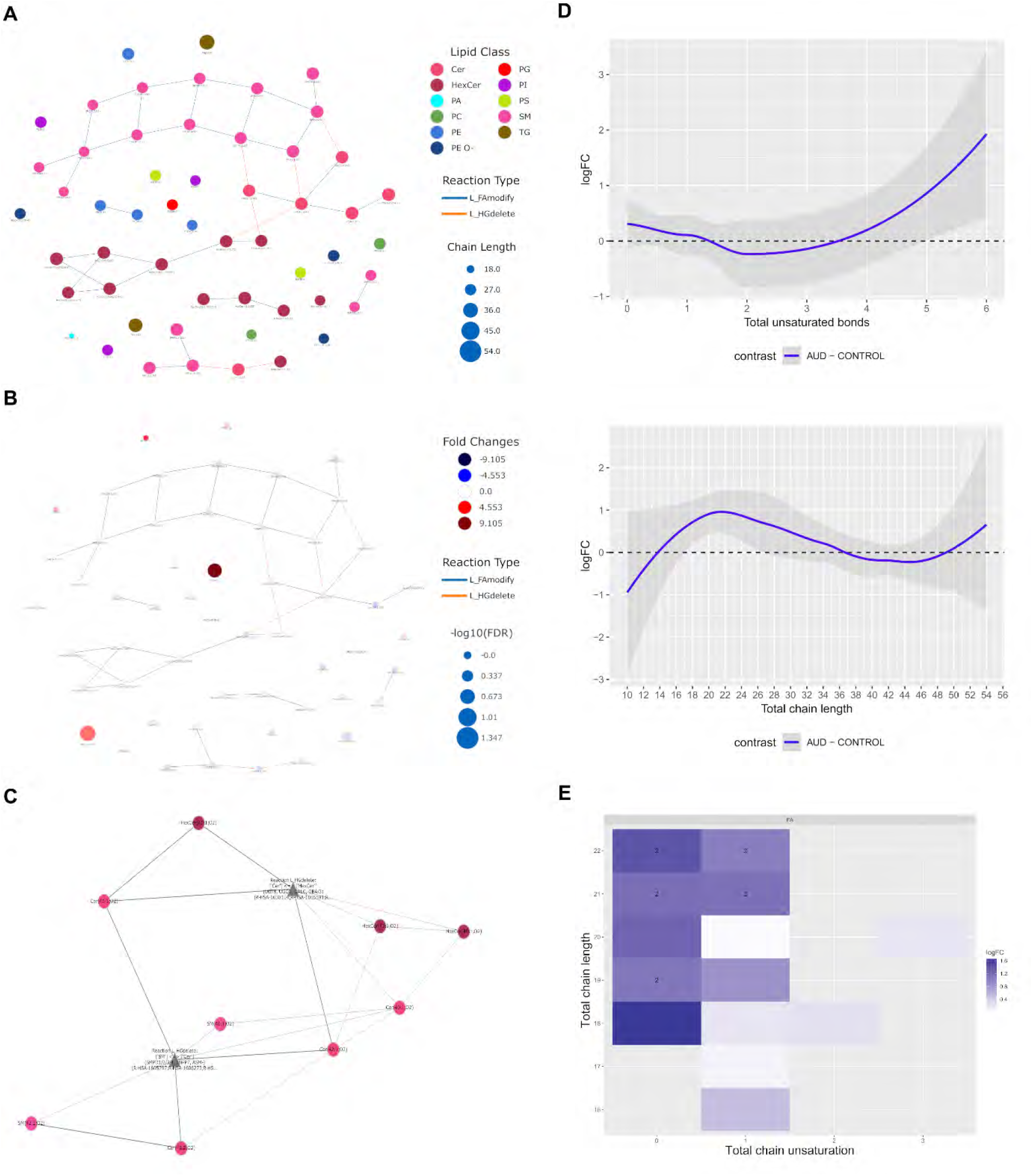
Functional lipid network analysis of urine extracellular vesicles (EVs) from male alcohol use disorder (AUD) patients and controls. **(A)** LINEX^2^-generated lipid reaction network based on LC–MS/MS data. Colored spherical nodes depict lipid classes. Edge colors indicate the type of reaction connecting nodes. **(B)** Global lipid network visualization with LINEX^2^ between AUD males and healthy individuals. Red spherical nodes represent lipids with a positive Fold Change (higher abundance in AUD), whereas blue nodes indicate a negative Fold Change (higher abundance in control). The spherical node sizes indicate the-log10 FDR corrected p-values of lipid species between AUD and control males (a larger node size represents a higher level of statistical significance). Edges are colored by correlation changes for lipids from AUD male patients to healthy individuals: negative to positive (significant correlation in both groups, <0 in AUD and >0 in control), positive to negative (significant correlation in both groups, >0 in AUD and <0 in control), significant to insignificant (significant correlation in AUD, insignificant in control), unchanged significant (significant in both groups, either both >0 or both <0), insignificant (uncorrelated in both groups), and insignificant to significant (insignificant in AUD, significant in control). **(C)** Enrichment subnetwork generated by LINEX^2^ based on the global network in AUD males. Spherical nodes represent lipid species, and triangular nodes represent reaction type. **(D)** Total unsaturated bonds and total chain length (number of carbons) in the lipid species in urine EVs of AUD male patients and controls. The LFC indicates the lipid differential abundance from AUD male patients to healthy individuals. Blue line with a positive LFC represents lipids with higher unsaturated bonds and chain length in AUD, whereas the blue line with a negative LFC indicates lipids with higher unsaturated bonds and chain length in controls. **(E)** Total unsaturated bonds and total chain length (number of carbons) in the subclass Fatty acids in urine EVs of AUD male patients and controls. The LFC indicates the lipid differential abundance from AUD male patients to healthy individuals.

Additionally, to expand the functional analysis (particularly regarding fatty acids not recognized by LINEX^2^), we performed a structural lipid analysis by examining chain length (number of carbons) and degree of unsaturation between AUD patients versus control. Figure 5D shows the total number of unsaturated bonds and overall chain length in lipid species present in urine EVs from AUD males and controls, demonstrating a higher abundance of lipids with chain lengths between 14 and 38 carbons in the AUD group. Figure 5E displays the total number of unsaturated bonds and total chain length in the fatty acid subclass, revealing increased acyl chain saturation and more lipid chains with 22–24 carbons in AUD males compared to controls.

## Discussion

We have previously demonstrated AUD-related lipidomic fingerprints in plasma EVs from AUD patients (Perpiñá-Clérigues et al., 2024), with the goal of identify lipid biomarkers for chronic heavy drinking diagnosis and prognosis. This study reveals that lipidome analysis of urine EVs from AUD males exhibits significant enrichment in fatty acyls and glycerophospholipids. Specifically, FA 22:0 emerges as a promising biomarker in urine EVs in AUD with recent ethanol intake, mirroring its abundance in plasma EVs of AUD males (Perpiñá-Clérigues et al., 2024). Notably, we report for the first time an increase in acyl chain saturation and very long-chain fatty acids (22-24 carbons) in AUD males, which may be linked to biological dysfunctions such inflammation, cancer, and metabolic disorders. It is worth noting that patients in our sample had been actively drinking in the weeks prior to inclusion and did not present with advanced liver disease. Thus, our results reflect alcohol-related metabolic changes, rather than consequences of end-stage organ damage.

Urine EVs have gained recognition as dynamic biomarkers for disease monitoring (Bajo-Santos et al., 2023; Hallal et al., 2024). While primarily associated with bladder and kidney diseases (Erdbrügger et al., 2021), they also show promise for cancer detection (Hallal et al., 2024) and neurological diseases (Sun et al., 2019; Wang et al., 2019). Compared to blood, urine offers distinct advantages, including non-invasive collection, larger sample volumes, and a less complex proteomic background (Harpole et al., 2016). Our lipidomic profiling identified 105 distinct lipid species in urine EVs, with prominent representation of unsaturated FAs, HexCer-AP, and sphingomyelins - contrasting with the 575 species detected in plasma EVs (enriched in TAG, PC, SM and unsaturated FA) (Perpiñá-Clérigues et al., 2024). This abundance may impact LC-MS/MS biomarker reliability.

Our results demonstrate that urine EVs from AUD males exhibit greater enrichment of fatty acyls and glycerophospholipid, alongside reduced sphingolipid. Ceramides and other sphingolipids are critical for membrane integrity and immune signaling, with implications in cancer, inflammation and neurodegeneration (Albeituni & Stiban, 2019; Jia et al., 2024). Furthermore, sphingolipids participate in EV biogenesis by modulating membrane curvature, influencing EV-mediated intercellular communication in both physiological and pathological contexts, including inflammatory and degenerative diseases (Verderio et al., 2018). Conversely, fatty acyls (e.g., FA 22:1, 19:0, 21:0 and 22:0) were elevated in AUD males, consistent with their roles in inflammation (Calder, 2011) and neurotransmitter release (Darios et al., 2007). To further explore potential AUD biomarkers in urine EVs, we directly compared the lipidomes of urine and plasma EVs. Strikingly, FA 22:0 (also named behenic acid or docosanoic acid) was significantly enriched in EVs from both biofluids in AUD males. A case–control study has reported elevated levels of this saturated fatty acid, behenic acid, in the postmortem frontal cortex of schizophrenia patients (Hamazaki et al., 2016), while hepatic lipidome analyses in non-alcoholic fatty liver disease patients linked FA 22:0 to liver fibrosis (Fridén et al., 2021). These findings collectively suggest that FA 22:0 in urine EVs could serve as a robust biomarker for AUD.

Integration of LINEX^2^-mediated lipid network enrichment enabled knowledge-based linkage of lipidomic and proteomic datasets, associating enzymatic functions with specific lipid species (Rose et al., 2023). Network analysis revealed dysregulation of Cer and SM metabolism in AUD males, involving enzymes such as ASM, ENPP7, SMPD family, and SGMS1. Our prior data on plasma EVs from AUD males similarly indicated alteration in these enzymes (Perpiñá-Clérigues et al., 2024), as did studies on chronic alcohol consumption (Barron et al., 2020). Other reports have documented aberrant sphingolipid levels (including Cer and SM) in alcohol-related liver disease (Longato et al., 2012) and high alcohol intake cohorts (Jaremek et al., 2013). The network analysis also highlighted dysregulation in Cer and HexCer metabolism, with key enzymes like UGT8 and UGCG implicated in glycosphingolipid pathways. For instance, UGCG (with synthesizes glucosylceramide, GlcCer) is overexpressed in fibrotic liver tissue and hepatic stellate cells, suggesting a role in fibrogenesis (Li et al., 2023). UGCG dysregulation has also been observed in hepatocellular carcinoma (Byrne et al., 2022), while UGT8 expression alterations were identified in postmortem hippocampal tissue from AUD patients (McClintick et al., 2013). These findings underscore the potential involvement of Cer/HexCer metabolism in alcohol-induced hepatic and neurological pathology.

Interestingly, lipid function is further influenced by acyl chain length and saturation, which critically affect membrane trafficking (Vanni et al., 2019). Here, we observed increased acyl chain saturation and elevated very long-chain FAs (22-24 carbons) in AUD males. Certain saturated fatty acids are known to promote inflammation (Berg et al., 2020; Mei et al., 2024), and a meta-analysis linked elevated circulating saturated fatty acids to higher cancer risk, particularly affecting breast, prostate, and colorectal cancers (Mei et al., 2024). Additionally, very long-chain fatty acids (>22 carbons) are implicated in metabolic disorders such as Zellweger syndrome and adrenoleukodystrophy (Kyselová et al., 2022).

While urine EV analysis and lipidomics represent cutting-edge approaches in biomarker research, this study has several limitations that warrant discussion. A critical challenge was the inclusion of sex-specific analyses. Although we initially aimed to compare both male and female participants, recruiting women with AUD proved exceptionally difficult. This reflects broader epidemiological trends, as alcohol consumption in women is often underreported due to social stigma and gender-specific drinking patterns (Erol & Karpyak, 2015; Gilbert et al., 2019). Consequently, the limited sample size of female AUD patients precluded meaningful gender comparisons, restricting our analysis to males. Methodological constraints further influenced our findings. Although we isolated urine EVs via ultracentrifugation—the current gold standard for EV purification—this technique is known to suffer from low yield and significant EV loss, potentially biasing downstream analyses (Théry et al., 2018). Isolated particles exhibited size and morphological characteristics consistent with exosomes, but due to the absence of specific exosomal markers, we conservatively classified them as generic EVs. Finally, lipidomic pathway analysis faced technical hurdles. The lack of standardized lipid nomenclature and inconsistent integration across computational tools complicates data interpretation. For example, the LINEX^2^ software, while useful for network enrichment, has inherent limitations: (1) it excludes free fatty acyls from functional analyses, and (2) its algorithm-generated results may introduce stochastic variability. These factors underscore the need for cautious interpretation of lipid pathway associations.

In conclusion, this study combines urine EV lipidomics as an innovative strategy to identify lipid biomarkers for AUD. We highlight FA 22:0 as a potential biomarker and reveal acyl chain saturation and very long-chain FAs as novel lipid signatures of AUD.

## Declarations

### Ethics approval and consent to participate

Human urine samples were used in accordance with the Declaration of Helsinki and were approved by the Ethics Committee of the University Hospital of Salamanca (PI2020/02432), and written informed consent was obtained from each participant.

### Supporting information

The following supporting information is available free of charge at ACS website.

Figure S1. Whole Western blots of CD9, CD63, CD81, and calnexin.

Table S1. Characteristics of the male individuals in the study who present chronic alcohol consumption.

Table S2. Abbreviation of different lipid subclasses.

Table S3. Classification by levels of all lipids present in samples.

Table S4. Lipids with significant differential abundance separated by logarithm of fold change (LFC).

### Data availability statement

The datasets generated and analyzed during the current study and programming scripts are available in the Zenodo repository, https://zenodo.org/records/15525237, and in a web platform: https://bioinfouvcipf.shinyapps.io/LipUrEVs-AUD/.

### Authors contributions

BMU analyzed the data; MP, CPC and FGG designed and supervised the bioinformatics analysis; MLAS, MRP, DPM, and MM obtained human urine samples and collected clinical data; SM isolated EVs from human urine and lipid fraction; CPC designed and implemented the web tool; BMU, CPC, MP and FGG wrote the manuscript; MP and SM and BMU designed the graphical abstract; BMU, CPC, MP and FGG helped in the interpretation of the results; BMU, CPC, SM, MM, MP and FGG writing-review and editing; MP and FGG conceived the work. All authors read and approved the final manuscript.

## Acknowledgments

The authors thank the Principe Felipe Research Center (CIPF) for providing access to the cluster, co-funded by European Regional Development Funds (FEDER) in the Valencian Community 2014-2020. The authors also thank the Genomics and Proteomics Unit at the University of Alicante, and the Electron Microscopy Service at the Príncipe Felipe Research Centre.

